# IL-1β disrupts blood-brain barrier development by inhibiting endothelial Wnt/β-catenin signaling

**DOI:** 10.1101/2023.12.04.569943

**Authors:** Audrey R. Fetsko, Dylan J. Sebo, Lilyana B. Budzynski, Alli Scharbarth, Michael R. Taylor

## Abstract

During neuroinflammation, the proinflammatory cytokine Interleukin-1β (IL-1β) impacts blood-brain barrier (BBB) function by disrupting brain endothelial tight junctions, promoting vascular permeability, and increasing transmigration of immune cells. Here, we examined the effects of Il-1β on the *in vivo* development of the BBB. We generated a doxycycline-inducible transgenic zebrafish model that drives secretion of Il-1β in the CNS. To validate the utility of our model, we showed Il-1β dose-dependent mortality, recruitment of neutrophils, and expansion of microglia. Using live imaging, we discovered that Il-1β causes a significant reduction in CNS angiogenesis and barriergenesis. To demonstrate specificity, we rescued the Il-1β induced phenotypes by targeting the zebrafish *il1r1* gene using CRISPR/Cas9. Mechanistically, we determined that Il-1β disrupts BBB development by decreasing Wnt/β-catenin transcriptional activation in brain endothelial cells. Given that several neurodevelopmental disorders are associated with inflammation, our findings support further investigation into the connections between proinflammatory cytokines, neuroinflammation, and neurovascular development.

## Introduction

The blood-brain barrier (BBB) performs a critical function in the central nervous system (CNS) by tightly regulating the passage of ions, molecules, and cells between the bloodstream and the brain.^1–5^ This barrier is primarily established by highly specialized brain endothelial cells that possess continuous tight junctions, efflux transporters, and low vesicular activity.^6–9^ Under normal physiological conditions, the BBB prohibits uncontrolled transcellular diffusion of hydrophilic molecules, harmful xenobiotics, many small molecule drugs, and most large biologics.^10, 11^ Thus, entry of essential proteins and water-soluble nutrients requires the expression of specific transporters by brain endothelial cells to enter the brain from circulation.^11, 12^ Furthermore, the BBB limits the transmigration of peripheral immune cells into the brain parenchyma under steady state conditions, although recent evidence indicates that the immune-privileged nature of the CNS is complex and not absolute.^13^ Conversely, CNS injuries, infections, and diseases induce proinflammatory signals that alter brain endothelial cell function and allow infiltration of immune cells across the BBB, resulting in neuroinflammation.^14–17^ While our understanding of the cellular and molecular mechanisms of neuroinflammation have advanced significantly over the past few decades, little is known about the consequences of proinflammatory stimuli during neurovascular development.

To establish a functional BBB, brain endothelial cells first begin to acquire essential barrier properties during the earliest stages of CNS angiogenesis in a process termed barriergenesis.^18, 19^ Eloquent studies in mice and zebrafish have revealed a cell autonomous requirement for Wnt/β-catenin signaling in brain endothelial cells to coordinate the Vegf-dependent migration of endothelial tip cells into the brain parenchyma and the acquisition of barrier properties.^20–23^ These developmental processes are mediated by neural progenitor cells that secrete Wnt7a and Wnt7b ligands, which signal through an endothelial cell receptor complex that includes Frizzled, Gpr124, Reck, and Lrp5/6.^22–29^ Once activated, Wnt/β-catenin signaling also promotes endothelial tight junction formation and expression of the Glucose transporter 1, Glut1 (a.k.a. Slc2a1), the first known marker of brain endothelial cell differentiation.^30–32^ Thus, Glut1 is frequently used as an early marker of Wnt/β-catenin dependent formation of the BBB.^19–21, 27, 28, 33–37^

Previous studies have also shown that endothelial Wnt/β-catenin signaling inhibits angiogenesis and normalizes tumor blood vessels in glioma models and is required for BBB integrity under neuropathological conditions such as ischemic stroke and glioblastoma.^38, 39^ These findings suggest that endothelial Wnt/β-catenin is required to stabilize BBB function and that aberrant Wnt/β-catenin may contribute to CNS disease pathology. Interestingly, Wnt/β-catenin signaling also exerts both anti-inflammatory and proinflammatory functions in a context dependent manner.^40^ For example, endothelial Wnt/β-catenin signaling reduces immune cell infiltration in both human multiple sclerosis and a mouse model of experimental autoimmune encephalomyelitis.^41^ Conversely, Wnt/β-catenin signaling stimulates NF-κB activity and promotes a proinflammatory phenotype in cultured endothelial cells treated with TNF-⍺.^42^ Despite the various interactions between endothelial Wnt/β-catenin signaling and inflammation, the impact of proinflammatory signals on BBB development has not been examined.

It is well-documented that CNS disease states that promote or exacerbate a neuroinflammatory response also cause BBB breakdown.^11, 12, 14, 15, 43^ These neuropathological processes are initiated by the expression of proinflammatory cytokines such as Interleukin-1β (IL-1β), Tumor Necrosis Factor alpha (TNF⍺), and Interleukin-6 (IL-6) among many others.^44–48^ In particular, IL-1β is a key mediator of the inflammatory response in the CNS and is associated with neurological conditions in which inflammation plays a prominent role. For example, IL-1β is upregulated in response to traumatic brain injury, ischemic stroke, and neurodegenerative disorders.^49–51^ Upon induction, IL-1β stimulates the expression of adhesion molecules on endothelial cells that facilitate the attachment and migration of immune cells across the BBB.^52^ Given the prominent role of IL-1β during neuroinflammation and the profound effects of IL-1β on brain endothelial cells, crosstalk between Wnt/β-catenin signaling and inflammatory pathways could potentially impact BBB development during embryogenesis. However, direct evidence indicating interactions between these signaling pathways in brain endothelial cells is lacking. Thus, our current study was designed to examine the impact of IL-1β expression during CNS angiogenesis and barriergenesis.

For these experiments, we utilized zebrafish (*Danio rerio*) as they provide an ideal model organism to: 1) visualize the in vivo development of the brain vasculature, 2) create transgenic lines to drive CNS-specific expression of Il-1β, and 3) analyze the effects of Il-1β on endothelial Wnt/β-catenin signaling during BBB development. Here, we generated a doxycycline-inducible transgenic system to promote the expression of Il-1β in the developing CNS. To establish the effectiveness of our model, we demonstrated a dose-dependent decrease in survival, brain infiltration of neutrophils, and the expansion of microglia/macrophages. Most importantly, we found a dose-dependent decrease in CNS angiogenesis and barriergenesis as indicated by reduced brain vasculature and decreased expression of barrier properties shown by our transgenic line driven by the zebrafish *glut1b* promoter.^19^ We also found a dose-dependent decrease in T-cell transcription factor (TCF) transgenic reporter activation in brain endothelial cells, indicating that Il-1β interferes with the Wnt/β-catenin pathway. To demonstrate that the effects of Il-1β were mediated through the Interleukin-1 receptor type 1 (Il1r1), we used CRIPSR/Cas9 to inactivate zebrafish Il1r1 and demonstrated rescue of the developmental phenotypes associated with Il-1β expression. In conclusion, our results reveal a previously unknown consequence of Il-1β on the developing brain vasculature and may provide important insights into the impact of neuroinflammation on BBB development.

## Results

### CNS-specific expression of Il-1β causes dose-dependent mortality and neuroinflammation

To drive the temporal and dose-dependent expression of Il-1β in the zebrafish CNS, we generated an inducible transgenic Tet-On model of neuroinflammation. This expression strategy utilizes two separate transgenic lines, a driver and a responder.^53^ For the driver line, we generated a Tol2 construct using the zebrafish *glial fibrillary acidic protein* (*gfap*) promoter upstream of the reverse tetracycline-controlled transactivator (rtTA) to produce pDestTol2CG2 *gfap:rtTA* (Figure 1A, left panel). The *gfap* promoter has previously been used to drive robust expression in the CNS during zebrafish development.^54–56^ For the reporter line, we generated another Tol2 construct using the tetracycline-responsive element (TRE) promoter upstream of a mature and secreted form of zebrafish Il-1β (*GSP-il1β^mat^*) to produce pDestTol2CmC2 *TRE:GSP-il1β^mat^* (Figure 1A, right panel). We previously demonstrated functionality of *GSP-il1β^mat^* by generating a systemic inflammation model using the Gal4-EcR/UAS system.^57, 58^ Both transgenic constructs were created with a corresponding transgenesis marker, *cmlc2:EGFP* (green myocardium) for the driver and *cmlc2:mCherry* (red myocardium) for the responder, for ease of transgene carrier identification (Figure 1A). Thus, in our Tet-On system, embryos generated from adult germline carriers of both transgenes secrete mature Il-1β within the CNS in a doxycycline (Dox)-dependent manner. For brevity throughout this study, we denote our model as “CNS/Il-1β”, which is comprised of the double transgenic lines *Tg(gfap:rtTA, cmlc2:EGFP)*, *Tg(TRE:GSP-il1β^mat^, cmlc2:mCherry)*.

**Figure 1.**
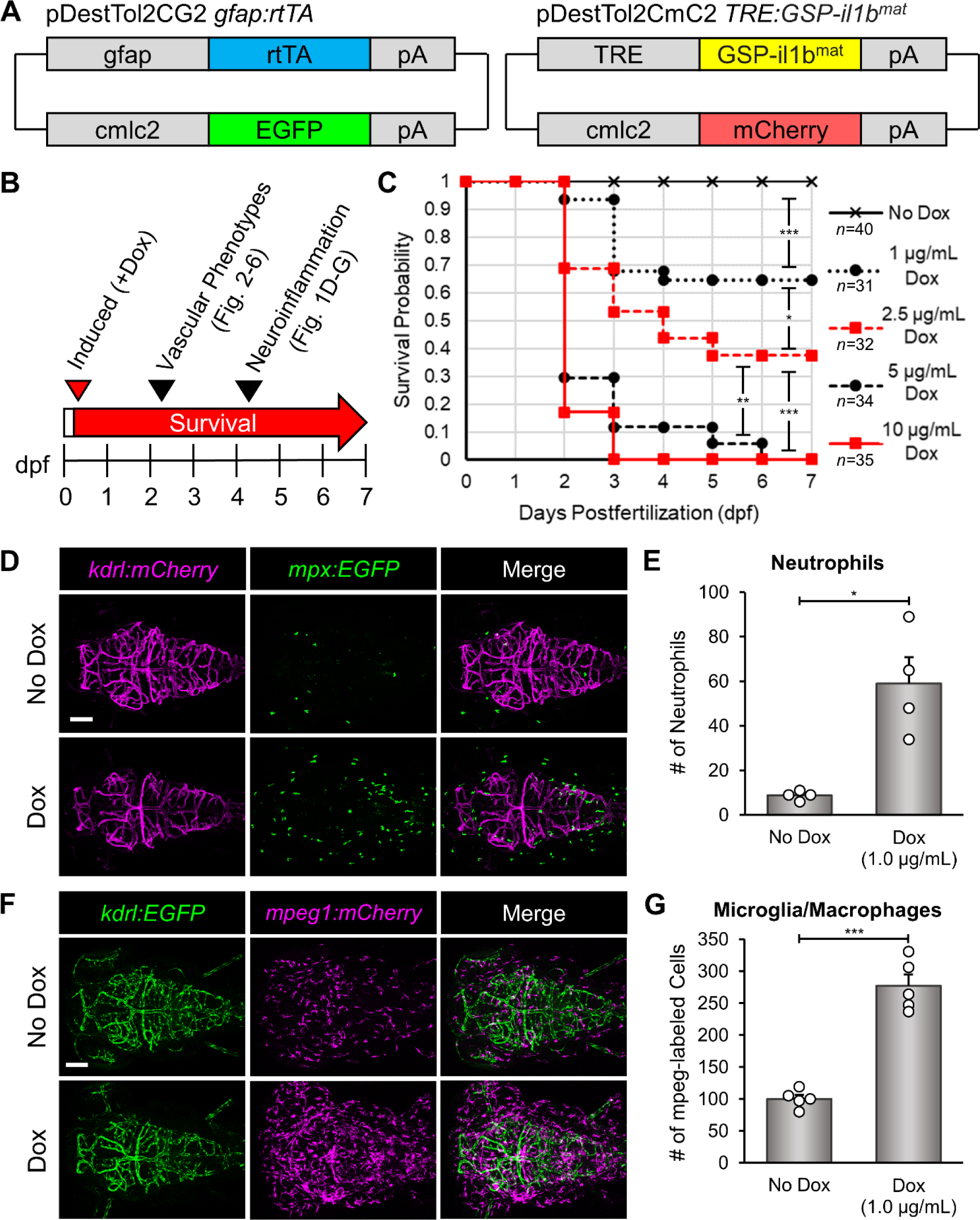
Dox-induced expression of Il-1β in the CNS promotes dose-dependent mortality and neuroinflammation in the transgenic CNS/Il-1β model. (**A**) Design of DNA constructs used to make the zebrafish transgenic lines *Tg(gfap:rtTA*, *cmlc2:EGFP)* and *Tg(TRE:GSP-il1b^mat^*, *cmlc2:mCherry*). The combination of these transgenic lines is designated as “CNS/Il-1β” to indicate doxycycline (Dox) inducible expression of Il-1β in the CNS. (**B**) Experimental timeline for all experiments. This graphic represents the timing of Dox induction and *in vivo* imaging. Dox (0-10 μg/mL) was added at approximately 6 hpf for all experiments. Survival was monitored daily through 7 dpf. Confocal imaging was performed at 4 dpf for neuroinflammation (Figure 1D and 1F) and at 2 dpf for imaging brain vasculature (Figures 2-6). (**C**) Kaplan-Meier analysis monitoring survival probability. CNS/Il-1β embryos were treated with Dox at 0, 1, 2.5, 5, or 10 μg/mL at 6 hpf, then monitored for survival every day until 7 dpf (*p < 0.05; **p < 0.01; ***p < 0.001; not all comparisons are shown). (**D**) Representative confocal microscopy images showing neutrophils (*mpx:EGFP*) and blood vessels (*kdrl:mCherry*) in the head. CNS/Il-1β embryos were untreated (No Dox) or treated (Dox; 1.0 μg/mL) at 6 hpf and then imaged at 4 dpf (dorsal view; anterior left). Scale bar is 100 μm. (**E**) Quantification of neutrophils (*mpx:EGFP*) in the heads of untreated (No Dox) or treated (Dox; 1.0 μg/mL) CNS/Il-1β larvae at 4 dpf (*n=*4). (**F**) Representative confocal microscopy images showing microglia/macrophages (*mpeg1:mCherry*) and blood vessels (*kdrl:EGFP*) in the head for context. CNS/Il-1β embryos at 6 hpf were either untreated (No Dox) or treated (Dox; 1.0 μg/mL), then imaged at 4 dpf (dorsal view; anterior left). Scale bar is 100 μm. (**G**) Quantification of microglia/macrophages (*mpeg1:mCherry*) in the heads of untreated (No Dox) or treated (Dox; 1.0 μg/mL) CNS/Il-1β larvae at 4 dpf (*n=*5). Error bars in E and G represent means ± SEM (*p < 0.05; ***p < 0.001).

To examine the effects of Il-1β expression in the CNS during embryonic development, we first monitored survival by performing a Kaplan-Meier analysis. For these experiments, Dox (0, 0.1, 5.0, and 10.0 µg/mL) was administered to CNS/Il-1β embryos at approximately 6 hours post-fertilization (hpf) and then monitored daily for survival until 7 days post-fertilization (dpf) (Figure 1B). As shown in Figure 1C, induction of Il-1β with Dox caused dose-dependent mortality. Treatment with 5 and 10 µg/mL Dox resulted in less than 20% of embryos surviving past 2-3 dpf. Approximately 40% of the 2.5 µg/mL Dox group and 60% of the 1.0 µg/mL Dox group survived to 7 dpf, and 100% of untreated (No Dox) embryos survived to 7 dpf. This rapid, dose-dependent mortality is comparable with the results of our previous studies using tebufenozide-induced Il-1β in a transgenic Gal4-EcR/UAS system designed to promote systemic inflammation.^57, 58^

Next, we examined the potential for our model to promote neuroinflammation. While the goal of this study was to examine the effects of Il-1β expression on brain vascular development, we also wanted to confirm that our CNS/Il-1β model promotes an inflammatory response similar to our Il-1β model of systemic inflammation.^57, 58^ In zebrafish, neutrophils (myeloperoxidase expressing granulocytic cells) begin to distribute throughout the embryo between 3-4 dpf, while yolk-derived macrophages begin to populate the brain parenchyma at about 35 hpf. These macrophages then transform into early microglia, the resident immune cells of the CNS, at approximately 55-60 hpf.^59, 60^ In our model, high concentrations of Dox (5.0-10.0 µg/mL) caused rapid mortality (Figure 1C), so we assessed neuroinflammation using a low concentration of Dox (1.0 µg/mL) to promote CNS inflammation without causing significant mortality by 4 dpf (Figure 1B). Under normal physiological conditions, neutrophils are absent from the CNS but transmigrate across the BBB during a neuroinflammatory response.^61^ Thus, to visualize neutrophils in the context of brain vasculature, we bred CNS/Il-1β to the *Tg*(*mpx:GFP)^uwm1^* (herein *mpx:GFP*) and *Tg(kdrl:HRAS-mCherry)^s896^* (herein *kdrl:mCherry*) transgenic lines and imaged embryos by confocal microscopy at 4 dpf. As expected, untreated embryos showed very few neutrophils in the head at 4 dpf. In contrast, Dox treatment caused a dramatic and significant increase of neutrophils indicative of neuroinflammation (Figure 1D and 1E). We also examined neutrophils in the brain at earlier developmental stages but did not observe any significant increases in response to Il-1β (Figure S1A and S1B).

We next quantified microglia/macrophages in the context of brain vasculature by imaging embryos derived from breeding CNS/Il-1β to the *Tg(mpeg1:mCherry)^gl23^* (herein *mpeg1:mCherry*) and *Tg(kdrl:EGFP)^s843^* (herein *kdrl:EGFP*) transgenic lines.^62,63^ Since the zebrafish *mpeg1* promoter drives expression in both microglia and macrophages, these specific cell types could not be distinguished for these experiments. Similarly to neutrophils, induction of Il-1β at early developmental stages did not show an increase in *mpeg1:mCherry* cells (Figure S1C and S1D). As expected for resident microglia, we observed *mpeg1:mCherry* cells in the heads of untreated embryos at 4 dpf. We also found an obvious and significant increase of *mpeg1:mCherry* cells in response to Dox treatment (Figure 1F and 1G). Together, the survival curves, infiltration of neutrophils, and increased number of microglia/macrophages demonstrate that our CNS/Il-1β model promotes an inducible, deleterious inflammatory response in the zebrafish CNS.

### Il-1β disrupts CNS angiogenesis in the transgenic CNS/Il-1β model prior to neuroinflammation

Neuroinflammation initiated by Il-1β is a well-known contributor to BBB dysfunction in the adult brain.^48^ However, the impact of Il-1β on CNS angiogenesis during embryonic development has not been examined. Therefore, we used our CNS/Il-1β model to evaluate the effects of Il-1β expression during early vascular development. While validating the inflammatory response in our model, we observed an obvious reduction in brain blood vessel density when treated with a low Dox concentration (Figure 1D and 1F; bottom left panels). In zebrafish, angiogenesis in the hindbrain begins at approximately 30 hpf when endothelial tip cells sprout into the brain parenchyma from the primordial hindbrain channels (PHBCs).^64, 65^ These emerging vessels migrate into the hindbrain and create the intracerebral central arteries (CtAs), which are mostly formed and connected to the basilar artery (BA) by 48 hpf.^66^ Therefore, to investigate the impact of Il-1β expression on the initial stages of CNS angiogenesis, we treated CNS/Il-1β, *kdrl:EGFP* embryos with varying doses of Dox at approximately 6 hpf and then imaged the brain vasculature by confocal microscopy at approximately 52 hpf (see experimental schematic in Figure 1B). As shown in Figure 2A, embryos treated with increasing Dox concentrations showed significantly reduced brain vasculature in comparison to untreated embryos. This effect was also dose dependent as shown for the survival curves (Figure 1C). The effects on CNS angiogenesis were quantified by counting the number of CtAs formed for each concentration of Dox (Figure 2B). Furthermore, we demonstrated that the reduction in angiogenesis was specific to the brain vasculature as shown by normal intersegmental vessel formation in the trunk of Dox-treated CNS/Il-1β embryos (Figure 2C and 2D).

**Figure 2.**
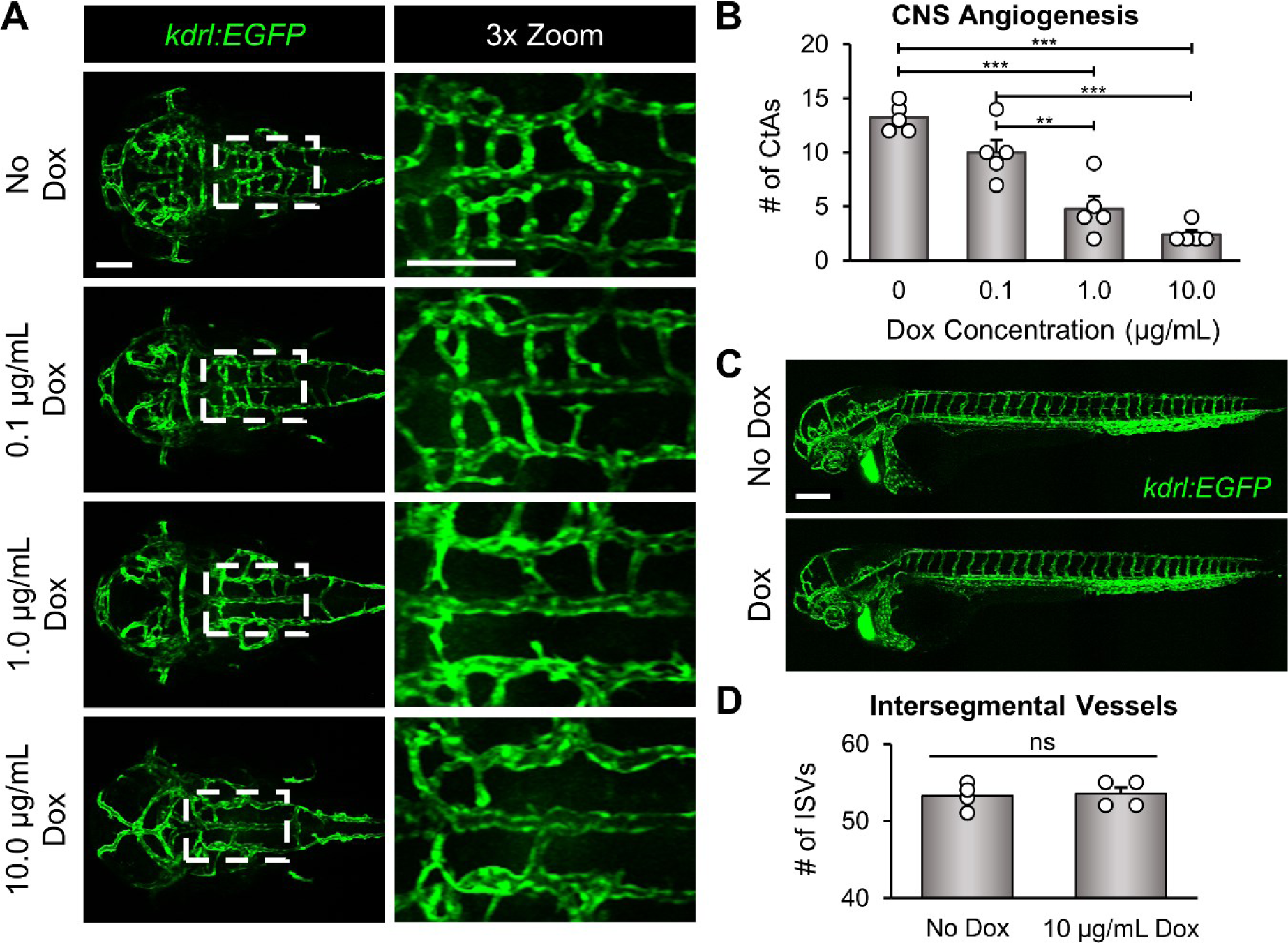
Il-1β expression disrupts CNS angiogenesis in the transgenic CNS/Il-1β model. (**A**) Representative confocal microscopy images showing dose-dependent effects of Dox on brain vascular development. CNS/Il-1β, *kdrl:EGFP* embryos were untreated (No Dox) or treated with Dox (0.1, 1.0, or 10.0 μg/mL) at 6 hpf and then imaged at 52 hpf (dorsal view; anterior left). Scale bars are 100 μm. (**B**) Quantification of the number of CtAs in CNS/Il-1β embryos treated with Dox (0, 0.1, 1.0, or 10.0 μg/mL) (*n*=5). (**C**) Representative confocal microscopy images showing whole embryo vasculature in CNS/Il-1β, *kdrl:EGFP* embryos at 2 dpf (lateral view; dorsal top; anterior left). CNS/Il-1β embryos were either untreated (No Dox) or treated (Dox; 10.0 μg/mL) at 6 hpf. Note the loss of CNS vasculature with Dox treatment but no effect on trunk vasculature with Dox treatment. Scale bar is 100 μm. (**D**) Quantification of the number of intersegmental vessels (ISVs) at 52 hpf in CNS/Il-1β embryos either untreated (No Dox) or treated (10.0 μg/mL Dox) (*n*=4). Error bars in B and D represent means ± SEM (**p < 0.01; ***p < 0.001; no label = not significant).

### CRISPR/Cas9 targeted deletion in the *il1r1* gene rescues mortality and CNS angiogenesis

We reasoned that the CNS angiogenesis defects were most likely due to Il-1β interactions with the Interleukin-1 receptor, type 1 (Il1r1) but conceded that these effects could also be caused by non-specific or undetermined mechanisms. Therefore, to determine if the Il-1β effects were mediated by the receptor, we used CRISPR/Cas9 to generate a conventional knockout of zebrafish Il1r1 by deleting the transmembrane domain to generate a null allele. Previous studies in mice have demonstrated that IL1R1 is required to elicit an immune response to IL-1β.^67, 68^ Furthermore, we previously showed that Il1β-driven systemic inflammation and associated phenotypes require Il1r1 and that morpholino or mosaic CRISPR/Cas9 knockdown of Il1r1 rescues these inflammatory effects.^58^ To generate a stable germline knockout of Il1r1, we generated two CRISPR/Cas9 ribonucleoprotein (RNP) complexes, cr1 and cr2, targeting exons 8 and 9 of *il1r1*, respectively (Figure 3A). CNS/Il-1β, *kdrl:EGFP* embryos were microinjected, raised to adulthood, and then genotyped by PCR. Next, embryos generated from heterozygous *il1r1*+/- adults were screened for rescue by treating with Dox and confirmed by PCR (Figure 3B). Embryos that survived Dox treatment were raised to adulthood and identified as *il1r1* deletion mutants (*il1r1*-/-). Adult *il1r1*-/- zebrafish were indistinguishable from wild type (Figure S2A). Similarly, *il1r1-/-* embryos showed no difference compared to wild type in the number of *mpeg1:mCherry*-positive in the brain at 30 and 52 hpf (Figure S2B and S2C). We also monitored survival by performing a Kaplan-Meier analysis. As shown in Figure 3C, *il1r1*-/- embryos survived Dox (10 µg/mL) treatment through day 7 whereas most Dox treated *il1r1*+/+ embryos did not survive past 2-3 dpf.

**Figure 3.**
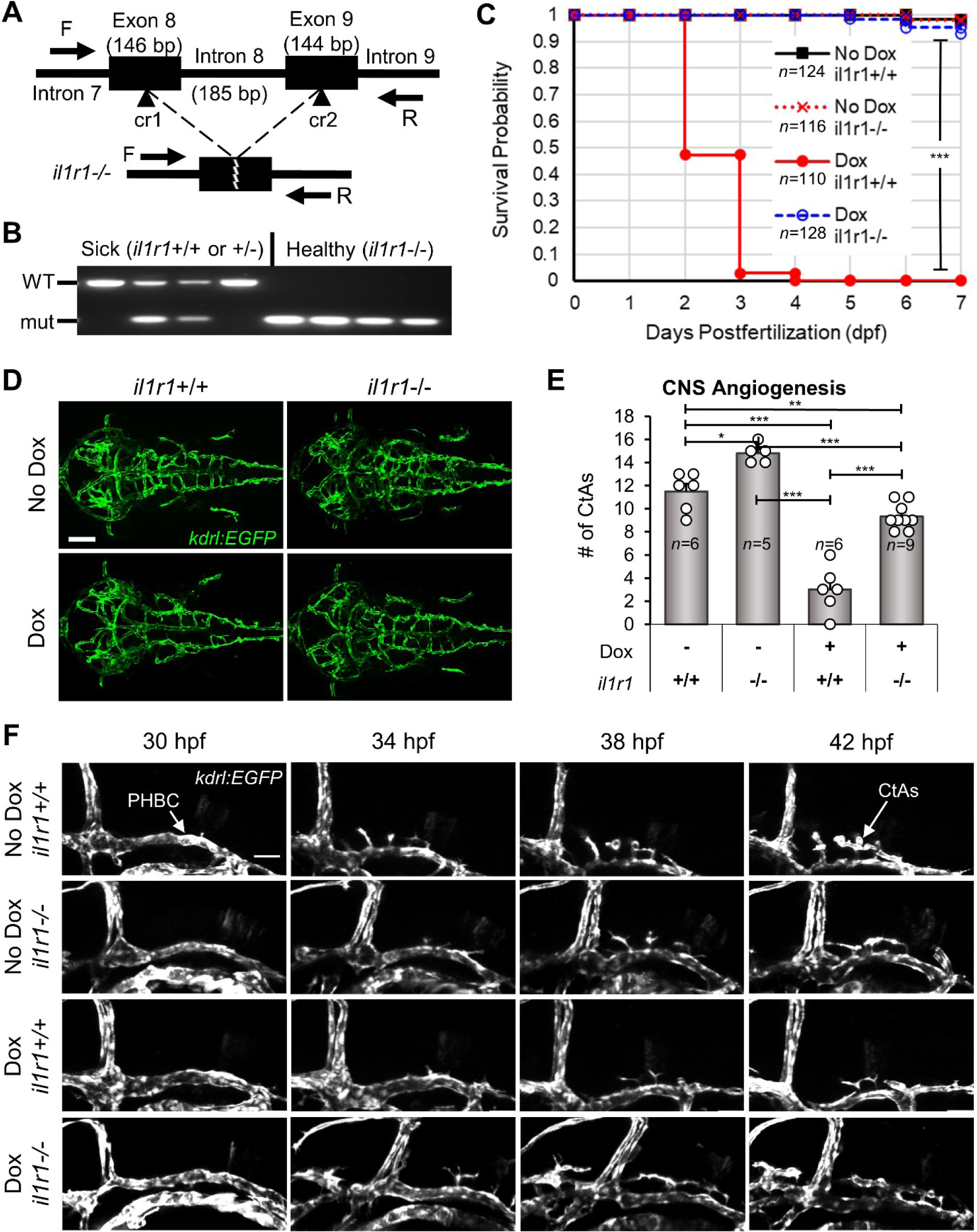
CRISPR/Cas9 *il1r1* mutants rescue Il-1β-induced mortality and CNS angiogenesis. (**A**) Schematic of the zebrafish *il1r1* gene showing the locations targeted by the CRISPR/Cas9 RNP complexes (cr1 and cr2) (top) and the resulting deletion of intron 8 (bottom). The exon and intron sizes in base pairs (bp) and the forward and reverse primers (F and R) used for genotyping are shown (not to scale). (**B**) PCR showing the genotyping of CRISPR/Cas9 deletion of *il1r1*. CNS/Il-1β embryos from heterozygous *il1r1*+/- adults were treated with Dox (10.0 μg/mL), selected as sick or healthy, and then genotyped by PCR. Note that all healthy embryos have the *il1r1*-/- deletion and all sick embryos carry a wild-type *il1r1* allele. (**C**) Kaplan-Meier analysis monitoring survival probability of *il1r1* mutants. CNS/Il-1β embryos either *il1r1*+/+ or *il1r1*-/- were untreated (No Dox) or treated (Dox; 10.0 μg/mL) at 6 hpf and then monitored for survival every day until 7 dpf (***p < 0.001; not all comparisons are shown). (**D**) Representative confocal microscopy images showing rescue of CNS angiogenesis in Dox-induced *il1r1* deletion mutants. CNS/Il-1β, *kdrl:EGFP* embryos either *il1r1*+/+ or *il1r1*-/- were untreated (No Dox) or treated (Dox 10 μg/mL) at 6 hpf and then imaged at 54 hpf (dorsal view; anterior left). Scale bars are 100 μm. (**E**) Quantification of the number of CtAs in wild-type *il1r1* (+/+) and mutant *il1r1* (-/-) embryos either treated with no Dox (-) or treated with 10.0 μg/mL Dox (+). Error bars represent means ± SEM (*p < 0.05; **p < 0.01; ***p < 0.001; no label = not significant). (**F**) Still frames from time-lapse confocal imaging of CNS/Il-1β, *kdrl:EGFP* embryos either *il1r1*+/+ or *il1r1*-/- without (No Dox) or with (10.0 μg/mL Dox) treatment at 6 hpf (lateral view; dorsal top; anterior left). Shown here are snapshots at 4-hour intervals over 12 hours of acquisition beginning at the onset of CNS angiogenesis (30 hpf). See Movies S1-4 for more detail.

To examine the effects on brain vascular development, we treated *il1r1*+/+ and *il1r1*-/- embryos with Dox (10 µg/mL) and then imaged the brain vasculature by confocal microscopy as described above. As with the survival analysis, we determined that deletion of *il1r1* rescued CNS angiogenesis in Dox treated embryos (Figure 3D). We quantified the number of CtAs and found a statistically significant rescue (Figure 3E). We also found that the number of CtAs was lower in the Dox-treated *il1r1*-/- embryos compared to untreated embryos, suggesting that Il-1β could potentially exert Il1r1-independent effects in the CNS as previously reported.^69, 70^ To visualize CNS angiogenesis in live embryos, we performed time-lapse resonant scanning confocal microscopy. Using Dox treated or untreated *il1r1*+/+ and *il1r1*-/- embryos, we captured z stacks at 20 min intervals from 30-42 hpf (Movies 1-4). Still frames from the time-lapse imaging are presented at 4-hour intervals (Figure 3F). Given that CRISPR/Cas9 targeted deletion of *il1r1* rescues both Il-1β induced mortality and defective CNS angiogenesis in our transgenic CNS/Il-1β model, we conclude that the deleterious effects of early embryonic expression of Il-1β are mediated through Il1r1.

### Il-1β disrupts *glut1b:mCherry* expression in brain endothelial cells during CNS angiogenesis

To determine whether expression of Il-1β in the CNS also impacts the acquisition of barrier properties (i.e. barriergenesis), we examined the induction of Glut1 in endothelial cells. As described above, Glut1 is the earliest known marker of barriergenesis and is frequently used as an indicator of BBB formation and function. Previous studies from our lab and others have demonstrated that zebrafish Glut1 is also an excellent marker of barriergenesis by either ⍺-Glut1 immunohistochemistry or transgenic reporter lines driven by the zebrafish *glut1b* promoter.^19–21, 34, 36^ Here, we bred CNS/Il-1β, *kdrl:EGFP* to our transgenic line *Tg(glut1b:mCherry)^sj1^* (herein *glut1b:mCherry*) and examined transgenic reporter expression in the PHBCs and CtAs following induction of Il-1β with a range of Dox (0, 0.1, 1.0, and 10.0 µg/mL). As shown in Figure 4A, Dox caused a dose-dependent decrease in *glut1b:mCherry* expression throughout the brain vasculature. We quantified *glut1b:mCherry* expression by measuring the mean fluorescence intensity of the mCherry signal within the hindbrain vasculature (see Materials and Methods for more details). The mean fluorescence intensity was significantly reduced in embryos treated with increasing concentrations of Dox in comparison to untreated embryos (Figure 4C). To further examine the reduction in mCherry signal in the hindbrain vasculature, the ratio of the length of *glut1b:mCherry*-positive vessels versus *kdrl:EGFP*-positive vessels was also quantified (Figure S3A). To control for the effects of Dox treatment alone, we imaged and quantified wild-type embryos (WT) in the transgenic *glut1b:mCherry*, *kdrl:EGFP* background without the CNS/Il-1β transgenes and found no significant impact on brain vascular development (Figure 4B, 4D, and 4E; Figure S3B).

**Figure 4.**
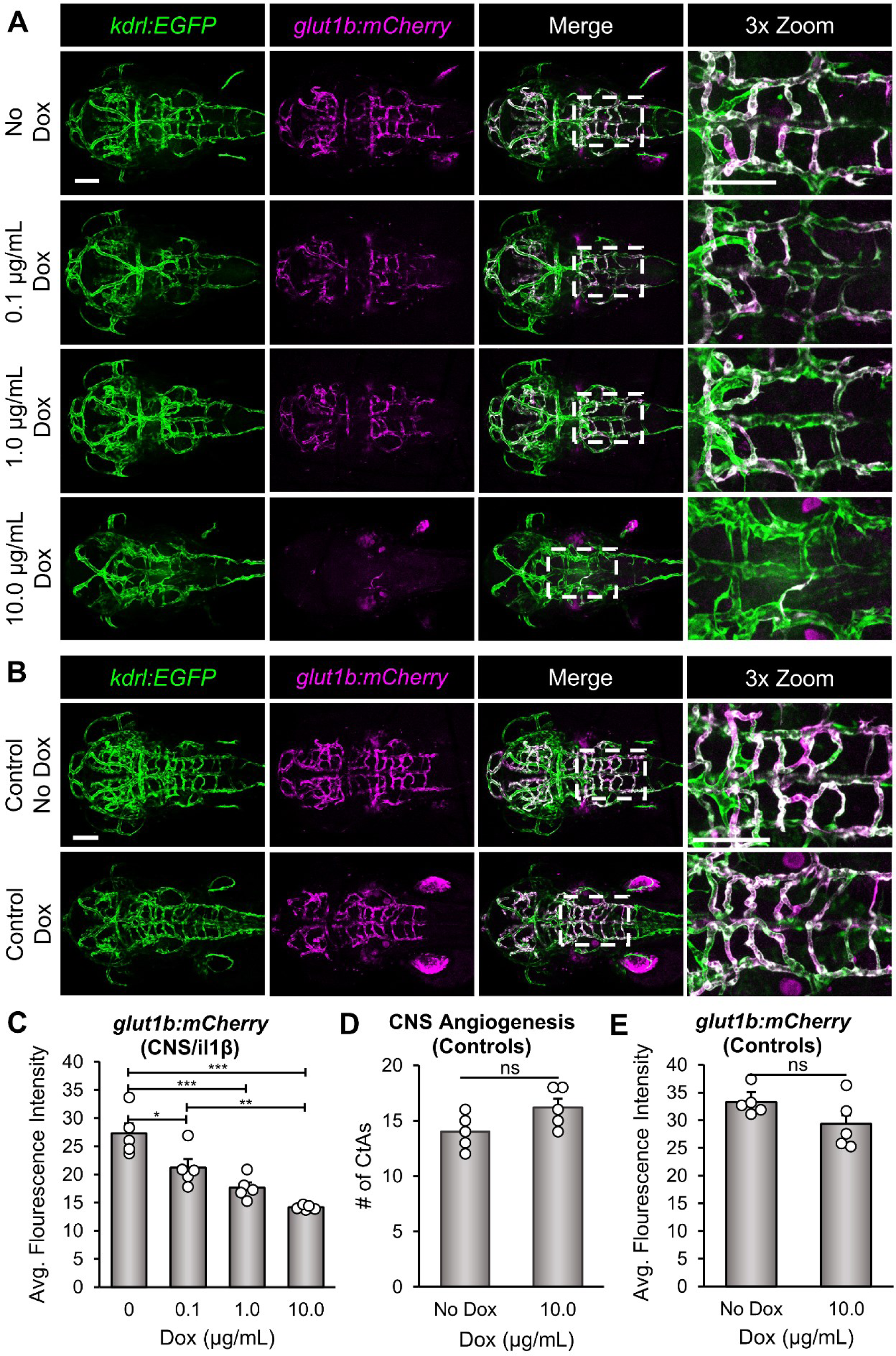
Il-1β disrupts *glut1b:mCherry* expression in brain endothelial cells during CNS angiogenesis. (**A**) Representative confocal microscopy images showing Dox-dependent effects on *glut1b:mCherry* expression. CNS/Il-1β, *kdrl:EGFP*, *glut1b:mCherry* embryos were untreated (No Dox) or treated with Dox (0.1, 1.0, or 10.0 μg/mL) at 6 hpf and then imaged at 52 hpf (dorsal view; anterior left). Scale bars are 100 μm. (**B**) Representative confocal microscopy images showing control *kdrl:EGFP*, *glut1b:mCherry* embryos (no CNS/Il-1β) at 2 dpf. Embryos were either untreated (No Dox) or treated with 10.0 μg/mL (Dox) at 6 hpf. Scale bars are 100 μm. (**C**) Quantification of the average *glut1b:mCherry* fluorescence intensity in the hindbrain vasculature of CNS/Il-1β, *kdrl:EGFP*, *glut1b:mCherry* embryos at 2 dpf (*n*=5). Note the dose-dependent decrease in *glut1b:mCherry* signal indicating a defect in barriergenesis. (**D,E**) Quantification of the number of CtAs (D) and average *glut1b:mCherry* fluorescence intensity (E) in the hindbrain vasculature of control *kdrl:EGFP*, *glut1b:mCherry* embryos (no CNS/Il-1β) without (No Dox) or with (10.0 μg/mL Dox) treatment (*n*=5). Note that Dox alone has no impact on barriergenesis. Error bars in C, D, and E represent means ± SEM (*p < 0.05; **p < 0.01; ***p < 0.001; ns or no label = not significant).

To examine permeability of the newly formed vessels, we performed microangiographic injections as previously described.^36^ For these experiments, we microinjected embryos at 48-52 hpf with DAPI and Texas Red dextran (10 kDa) and then imaged and quantified tracer leakage into the brain parenchyma adjacent to the PHBCs and in the tail region by confocal microscopy 30 minutes after injection. These experiments were performed on untreated (No Dox) and treated (1.0 μg/mL Dox) embryos. We were unable to perform these experiments on embryos treated with 10.0 μg/mL Dox due to significant morbidity and reduced circulation. Our results did not show a statistically significant difference between the No Dox and 1.0 μg/mL Dox embryos using the Texas Red dextran (10 kDa) tracer, indicating that Il-1β does not increase (or decrease) leakage at this developmental stage (Figure S4A and S4B). In addition, we did not observe DAPI stained nuclei in brain parenchymal cells in either the No Dox or 1.0 μg/mL Dox embryos as reported in the Tam et al study (Figure S4A; left panels). However, we did observe brain endothelial cell nuclei staining and extravascular nuclei staining in the tail region in both the No Dox and 1.0 μg/mL Dox embryos (Figure S4A). The low level of leakage at 2 dpf in both the treated and untreated embryos is consistent with recent studies in zebrafish suggesting that BBB tightness does not occur until 3-5 dpf.^71, 72^

In line with the CRISPR/Cas9 rescue of survival and CNS angiogenesis (Figure 3), we predicted that knockdown of Il1r1 would also rescue the expression of *glut1b:mCherry* in brain endothelial cells. For these experiments, we used CRISPR/Cas9 to generate mosaic knockdown of Il1r1 in CNS/Il-1β, *kdrl:EGFP, glut1b:mCherry* transgenic embryos. Our previous studies demonstrated the feasibility of this knockdown strategy by rescuing mortality caused by Il-1β induced systemic inflammation.^58^ The advantage of this experimental paradigm is that it bypasses the need to generate adult *il1r1*-/- in multiple transgenic backgrounds, eliminating additional rounds of breeding and selection. Here, CNS/Il-1β, *kdrl:EGFP, glut1b:mCherry* single-cell embryos were co-injected with CRISPR/Cas9 RNP complexes, cr1 and cr2, to generate “*il1r1* crispants” (see Figure 3A). As shown in Figure 5A, *il1r1* crispants showed normal CNS angiogenesis and barriergenesis in the absence of Dox and substantial rescue of Il-1β induced phenotypes in the presence of Dox. To quantify these observations, we counted the number of CtAs and measured the *glut1b:mCherry* fluorescence intensities. We found that Dox treated *il1r1* crispants showed a statistically significant increase in CtAs and *glut1b:mCherry* expression, although the rescue was not absolute (Figure 5B and 5C; Figure S3C). We previously showed that *il1r1* mRNA is present in zebrafish embryos at 0 dpf; therefore, we suspect that this nominal effect is due to maternally derived *il1r1* transcripts.^58^

**Figure 5.**
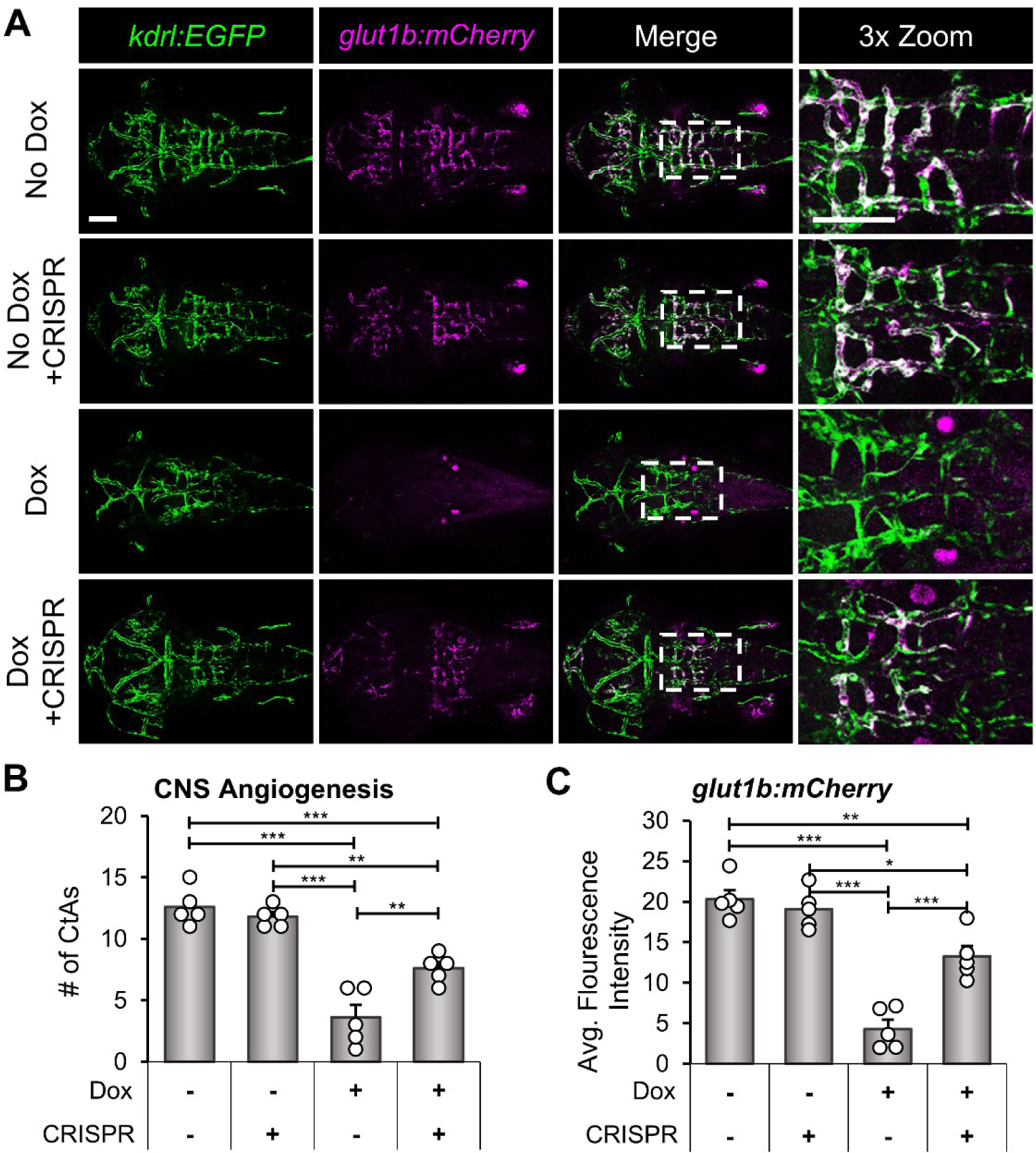
*il1r1* crispants rescue *glut1b:mCherry* expression in brain endothelial cells during CNS angiogenesis. (**A**) Representative confocal microscopy images showing rescue of *glut1b:mCherry* expression in *il1r1* crispants. CNS/Il-1β, *kdrl:EGFP*, *glut1b:mCherry* embryos were injected with CRISPR/Cas9 RNP complexes (cr1 and cr2) at the one-cell stage. Control embryos and *il1r1* crispants were either untreated (No Dox) or treated (10.0 μg/mL Dox) at 6 hpf and then imaged at 52 hpf (dorsal view; anterior left). Scale bars are 100 μm. (**B,C**) Quantification of the number of CtAs (B) and average *glut1b:mCherry* fluorescence intensity (C) in the hindbrain vasculature of control (CRISPR -) and *il1r1* crispants (CRISPR +) either untreated (Dox -) or treated with 10.0 μg/mL Dox (Dox +) (*n*=5 for each condition). Error bars in B and C represent means ± SEM (*p < 0.05; **p < 0.01; ***p < 0.001; no label = not significant).

### Il-1β reduces Wnt/β-catenin dependent TCF transcriptional activation in brain endothelial cells

Endothelial Wnt/β-catenin signaling is known to play a central role in both CNS angiogenesis and barriergenesis.^18, 22^ Given that induction of Il-1β in the CNS caused a significant decrease in CtA formation and *glut1b:mCherry* expression, we predicted that Wnt/β- catenin signaling may be perturbed in brain endothelial cells as a result of Il-1β expression during neurovascular development. Furthermore, we recognized that the Il-1β induced vascular phenotypes are very similar to that of zebrafish Wnt co-receptor mutants, *gpr124* and *reck*, that disturb endothelial Wnt/β-catenin.^19–21^ For example, both *gpr124* and *reck* mutants showed a significant reduction in the number of CtAs, loss of Glut1 expression as shown by IHC or the *glut1b:mCherry* transgenic reporter, and decreased Wnt/β-catenin transgenic reporter activation.^19–21^ Therefore, we examined activation of *Tg(7xTCF-Xla.Sia:NLS-mCherry)^ia5^* (herein *TCF:mCherry*), a transgenic Wnt/β-catenin transcriptional reporter line with a nuclear localization signal, to determine the level of Wnt/β-catenin transcriptional activation in Dox-treated CNS/Il-1β embryos.^73^ Here, we bred CNS/Il-1β, *kdrl:EGFP* to *TCF:mCherry* and examined transgenic reporter expression in the PHBCs and CtAs following induction of Il-1β with a range of Dox (0, 0.1, 1.0, and 10.0 µg/mL). As shown in Figure 6A, the highest concentration of Dox (10.0 µg/mL) resulted in the loss of *TCF:mCherry*-positive nuclei in the hindbrain vasculature, whereas 1.0 µg/mL and 0.1 µg/mL Dox showed no significant change. These observations were quantified by counting the number of *TCF:mCherry*-positive nuclei per length of blood vessel in the hindbrain vasculature (Figure 6B). To control for the effects of Dox treatment alone, we imaged and quantified wild-type embryos (WT) in the transgenic *kdrl:EGFP*, *TCF:mCherry* background without CNS/Il-1β and found no significant impact on endothelial Wnt/β-catenin activity (Figure 6C and 6D). Together with the significant reduction in CNS angiogenesis and barriergenesis, these data indicate that CNS expression of Il-1β disrupts Wnt/β-catenin signaling in brain endothelial cells during neurovascular development.

**Figure 6.**
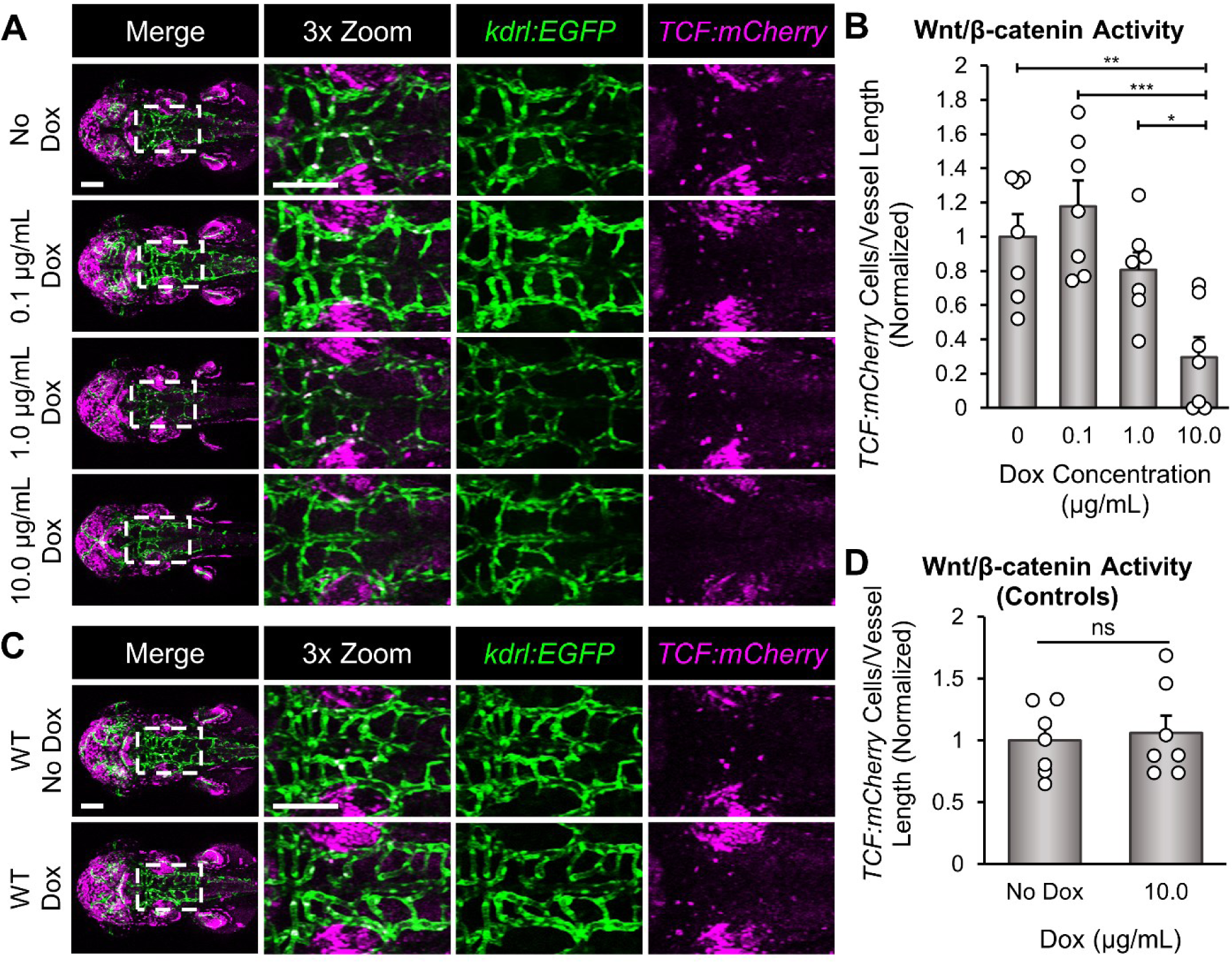
Il-1β reduces Wnt/β-catenin transcriptional activation in brain endothelial cells. (**A**) Representative confocal microscopy images showing Wnt/β-catenin transcriptional activity. CNS/Il-1β, *kdrl:EGFP*, *TCF:mCherry* embryos were untreated (No Dox) or treated with Dox (0.1, 1.0, or 10.0 μg/mL) at 6 hpf and then imaged at 52 hpf (dorsal view; anterior left). Scale bars are 100 μm. (**B**) Quantification of *TCF:mCherry*-positive endothelial cells in the hindbrain vasculature (*n*=7). Values are the number of *TCF:mCherry*-positive cells in *kdrl:EGFP*-positive blood vessels divided by the length of the hindbrain vasculature in each embryo. All values were normalized to the average of the No Dox group. (**C**) Representative confocal microscopy images showing control *kdrl:EGFP*, *TCF:mCherry* embryos (no CNS/Il-1β) at 2 dpf (dorsal view; anterior left). Embryos were either untreated (No Dox) or treated with 10.0 μg/mL (Dox) at 6 hpf. (**D**) Quantification of *TCF:mCherry*-positive endothelial cells in the hindbrain vasculature (*n*=7). Values are the number of *TCF:mCherry*-positive cells in *kdrl:EGFP*-positive blood vessels divided by the length of the hindbrain vasculature in each embryo. All values were normalized to the average of the No Dox group. Scale bars for A and C are 100 μm. Error bars in B and D represent means ± SEM (*p < 0.05; **p < 0.01; ***p < 0.001; ns or not labeled = not significant).

## Discussion

In this study, we discovered that CNS-specific expression of Il-1β interferes with BBB development by disrupting Wnt/β-catenin signaling in brain endothelial cells. We developed a doxycycline-inducible transgenic zebrafish model, CNS/Il-1β, that promotes secretion of mature Il-1β by radial glial cells, the primary neural progenitor cells during embryonic development.^74^ To demonstrate the utility of our model, we showed that Il-1β causes dose-dependent mortality, recruitment of neutrophils, and expansion of microglia/macrophages. As Il-1β is known to promote neuroinflammation and disrupt BBB function, the goal of our study was to examine the effects of Il-1β on BBB development.^48^ Here, we showed that CNS expression of Il-1β during neurovascular development causes a significant reduction in CNS angiogenesis and barriergenesis as indicated by decreased CtA formation and *glut1b:mCherry* expression, respectively. To demonstrate specificity of these effects, we rescued the Il-1β induced phenotypes using CRISPR/Cas9 against the gene encoding the zebrafish Interleukin-1 receptor, type 1 (Il1r1).^58^ In addition, we recognized that the brain vascular phenotypes were reminiscent of zebrafish *gpr124* and *reck* mutants.^19–21^ Since both Gpr124 and Reck function as receptor cofactors for Wnt ligands Wnt7a/Wnt7b and Wnt/β-catenin signaling is essential for CNS angiogenesis and barriergenesis, we reasoned that Il-1β likely interferes with endothelial Wnt/β- catenin signaling. Indeed, we found that Wnt/β-catenin transcriptional activation in brain endothelial cells is significantly decreased following the induction of Il-1β.

Since Il-1β appears to obstruct endothelial Wnt/β-catenin transcriptional activation via Il1r1, we predict substantial crosstalk between the nuclear factor-kappa B (NF-κB) and the Wnt/β-catenin signaling pathways during BBB development. When a neuroinflammatory response is triggered, Il-1β (as well as other proinflammatory cytokines) is produced by multiple cell types within the CNS. Once secreted, Il-1β binds to Il1r1 and recruits Il1rap to form a functional receptor complex and then regulates gene transcription via the NF-κB pathway.^75^ Comparably, activation of the Wnt/β-catenin pathway regulates gene transcription via nuclear translocation of β-catenin, which interacts with TCF/LEF to form an active transcriptional complex.^76–78^ These two pathways can reciprocally influence each other’s activities both positively and negatively, with the specifics of these interactions being highly dependent on the cellular, developmental, and disease context.^40^ For example, activation of the NF-κB pathway can interfere with β-catenin activity by inhibiting its translocation to the nucleus.^79^ In addition, NF-κB signaling can also indirectly obstruct Wnt/β-catenin signaling by inducing the expression of genes that promote the proteasomal degradation of β-catenin.^80^ Similarly, binding of Il-1β to the Il1r1/Il1rap receptor complex can activate the Mitogen-Activated Protein Kinase (MAPK) signaling pathway, and MAPK, as well as related protein kinases, can modulate β-catenin stability.^81–83^ Thus, the inhibitory effects of Il-1β on Wnt/β-catenin signaling are an example of the complex interactions between inflammatory and developmental signaling pathways that regulate cellular processes in health and disease. Furthermore, this intersection of pathways may be a newly discovered interaction in brain endothelial cells as we are not aware of any current studies that demonstrate this effect.

While we showed that Il1r1-dependent disruption of endothelial Wnt/β-catenin signaling impedes normal BBB development, we must also consider that Il-1β could affect other cell types in the CNS. For example, non-endothelial activation of the NF-κB and MAPK pathways could indirectly impact endothelial Wnt/β-catenin signaling by promoting the expression of inhibitory factors such as Dickkopf 1 (Dkk1) and Wnt inhibitory factor 1 (Wif1).^40^ Secretion of these antagonists could potentially inhibit the Wnt receptor complex on brain endothelial cells and prevent activation of Wnt/β-catenin signaling, thus inhibiting BBB development. Along these lines, secretion of Wnt inhibitory factors from Wnt-medulloblastoma disrupts the BBB phenotype in the tumor vasculature by blocking endothelial Wnt/β-catenin signaling.^84^ However, this indirect mechanism appears unlikely in our model. Given that CNS angiogenesis and barriergenesis begins early in development (30 hpf), the direct effects of Il-1β on endothelial Wnt/β-catenin signaling seem more plausible. Our previous studies showed unrestricted expression of zebrafish *il1r1* in the embryonic brain by *in situ* hybridization, thus endothelial Il1r1 expression seems likely.^58^ In the adult mouse brain, IL1R1 is primarily expressed by endothelial, ependymal, and choroid plexus cells and dentate gyrus neurons, scarcely expressed by astrocytes, and not expressed by microglia or perivascular macrophages.^85^ In contrast, the embryonic zebrafish brain is devoid of many of these adult cell types that respond to Il-1β as the choroid plexus does not form until 3 dpf, microglia differentiate after 2 dpf, and anatomical structures such as the hippocampus and dentate gyrus are not present.^86, 87^ Given this evidence, we predict that endothelial Il1r1 manifests the direct effects of Il-1β expression in the CNS. Regardless of the precise mechanism(s), our study provides the first direct observation that Il-1β disrupts BBB development by interfering with endothelial Wnt/β-catenin signaling.

In conclusion, Il-1β is a potent proinflammatory cytokine that modulates various physiological and pathological processes within the CNS. During a neuroinflammatory response, Il-1β is produced by multiple cell types and manifests its actions via binding to the Il1r1/Il1rap receptor complex. Brain endothelial cells that form the BBB are particularly affected by Il-1β. For example, Il-1β is known to disrupt tight junctions in brain endothelial cells which compromises BBB integrity and causes vascular permeability. In addition, Il-1β activates brain endothelial cells leading to the expression of adhesion molecules and the secretion of chemokines. Together, these effects result in the transmigration of immune cells across the BBB, exacerbating neuroinflammation. While the detrimental effects on the BBB have been well documented in the adult brain, little is known about the developmental consequences of Il-1β on brain endothelial cells. In this study, we discovered that Il-1β compromises BBB development during embryogenesis. These findings raise important questions about the effects of Il-1β and other cytokines during fetal development as the brain is particularly vulnerable *in utero*. Several neurodevelopmental disorders have been linked to early maternal immune activation (MIA) and inflammation, including autism spectrum disorder (ASD), schizophrenia, cerebral palsy, and attention-deficit/hyperactivity disorder (ADHD).^88, 89^ Given our novel findings of Il-1β induced cerebrovascular defects during early development, further investigation into the links between proinflammatory cytokines, neuroinflammation, and neurodevelopmental disorders is warranted.

### Limitations of the study

In this study, we generated doxycycline inducible transgenic zebrafish lines to examine the *in vivo* effects of Il-1β on BBB development. We are unable to determine the level of Il-1β protein expression due to the lack of antibodies that cross react with the zebrafish protein. We have addressed this limitation by showing dose-dependent inflammatory responses using varying concentrations of doxycycline to induce Il-1β expression. We also demonstrate that doxycycline alone had no impact on the phenotypes examined. Additionally, the CRISPR/Cas9 generated deletion of zebrafish *il1r1* was a conventional knockout and not a conditional knockout. Thus, cell type-specific effects of Il1r1 deletion were not examined in this study.

## Supporting information

Movie 1: Control No Dox 12h Time lapse

Movie 2: il1r1-/- No Dox 12h Time lapse

Movie 3: Control 10 Dox 12h Timelapse

Movie 4: il1r1-/- 10 Dox 12h Time lapse

## Author Contributions

ARF conducted the experiments, collected, assembled, analyzed the data, and wrote the manuscript. DJS provided experimental support and helped in data analysis. LBB and AS provided experimental support. MRT conceived the study, designed experiments, interpreted data, and wrote the manuscript. All authors contributed to the article and approved the submitted version.

## Acknowledgements

We thank Dr. Junsu Kang (UW-Madison) for providing the *Tg(7xTCF-Xla.Sia:NLS-mCherry)^ia5^* transgenic line, Dr. Owen Tamplin (UW-Madison) for providing the *Tg(mpeg1:mCherry)^gl23^* transgenic line, Dr. Anna Huttenlocher (UW-Madison) for providing the *Tg(mpx:EGFP)^uwm1^* transgenic line, and Dr. Jan Huisken (Morgridge Institute for Research and UW-Madison) for providing the *Tg(kdrl:HRAS-mCherry)^s896^* transgenic line. We also wish to thank Randall Kopielski for assistance with zebrafish husbandry and animal facility care and maintenance. This project was funded by NIH RO1NS116043. ARF was funded by the UW-Madison Biotechnology Training Program, NIH T32GM008349 and T32GM135066.

## Materials and Methods

### RESOURCE AVAILABILITY

#### Lead contact

Further information and requests for resources and reagents should be directed to and will be fulfilled by the lead contact, Michael R. Taylor.

#### Materials availability

Plasmids and zebrafish lines from this study are available from the lead contact upon request.

#### Data and code availability

All data reported in this paper will be shared by the lead contact upon request.

### EXPERIMENTAL MODEL DETAILS

#### Zebrafish husbandry and experimental lines

Zebrafish were maintained and bred using standard practices.^90^ Embryos and larvae were maintained at 28.5°C in egg water (0.03% Instant Ocean in reverse osmosis water). For imaging, 0.003% phenylthiourea (PTU) (TCI Chemicals) was used to inhibit melanin production. The transgenic zebrafish lines *Tg(gfap:rtTA)* and *Tg(TRE:GSP-Il1β^mat^*) were generated as described below, and the line *Tg(glut1b:mCherry)^sj1^* was previously generated in our lab.^19^

The transgenic zebrafish lines *Tg(kdrl:HRAS-mCherry)^s896^* and *Tg(kdrl:EGFP)^s843^* were provided by Dr. Jan Huisken (Morgridge Institute for Research and UW-Madison).^63, 91^ *Tg(mpx:EGFP)^uwm1^* was a gift from Dr. Anna Huttenlocher (UW-Madison).^92^ *Tg(mpeg1:mCherry)^gl23^* was provided by Dr. Owen Tamplin (UW-Madison).^62^ The line *Tg(7xTCF-Xla.Sia:NLS-mCherry)^ia5^* was provided by Dr. Junsu Kang (UW-Madison).^73^

All experiments were performed in accordance with the University of Wisconsin-Madison Institutional Animal Care and Use Committee (protocol number M005020).

#### Generation and Dox-induction of the *Tg(gfap:rtTA)*, *Tg(TRE:GSP-Il1β^mat^*) transgenic system (i.e. CNS/Il-1β)

Plasmids were constructed using Gateway cloning and components of the Tol2kit.^93, 94^ The plasmids *pME-rtTA* and *p3E-TRE* were made by inserting the *Tet-On* fragments from the Tet-On system (Clontech) in the appropriate Tol2 entry vectors. For *p5E-gfap*, the zebrafish *gfap* promoter was released from the *pGFAP-EGFP* vector and inserted into the 5’ entry clone *p5E-MCS*.^54^ The *pME:GSP-il1β^mat^* vector was constructed in our lab previously, as described by Lanham et al.^57^

To make the pDestTol2CG2 *gfap:rtTA* construct, *p5E:gfap*, *pME:rtTA*, *p3E:pA*, and the destination vector pDestTol2CG2 were recombined using LR Clonase II Plus (Invitrogen). To make the pDestTol2CmC2 *TRE:GSP-il1β^mat^* construct, *p5E:TRE*, *pME:GSP-il1β^mat^*, *p3E:pA*, and the destination vector pDestTol2CmC2 were recombined in the same manner.

To generate the transgenic lines *Tg(gfap:rtTA, cmlc2:EGFP)* and *Tg(TRE:GSP-il1β^mat^, cmlc2:mCherry)*, the Tol2 constructs were co-injected into single-cell embryos either individually or in combination (50–100 pg total plasmid DNA) together with 20 pg of *in vitro* transcribed *Tol2* transposase mRNA in a final volume of 1–2 nanoliters. Embryos with strong transient expression of the transgenesis markers were raised to adulthood and screened for germline transmission of the transgenes.

To induce expression of Il-1β, double transgenic *gfap:rtTA*; *TRE:GSP-il1β^mat^* embryos were treated with 0.1 – 10 μg/mL doxycycline (Dox) at approximately 6 hpf. Dox (RPI) was stored at −20°C as 10 mg/mL stocks in water and added to PTU/egg water or egg water without PTU for survival analysis at the desired concentration. All experiments were performed on embryos that were heterozygous for both the *gfap:rtTA* and *TRE:GSP-il1β^mat^* transgenes.

### METHOD DETAILS

#### Confocal laser scanning microscopy

For live imaging, zebrafish embryos and larvae were anesthetized in 0.02% tricaine (Western Chemical Inc.) and imbedded in 1.2% low melting point agarose (Invitrogen) in egg water with 0.003% PTU in a 35 mm glass-bottom dish, number 1.5 (MatTek). Confocal microscopy was performed using a Nikon Eclipse Ti microscope equipped with a Nikon A1R. For images of whole embryos and larvae, large images (4 x 1 mm) were captured and stitched together with a 15% overlap. For timelapse imaging, resonant scanning was used to acquire z-stacks at 20 min intervals for 12 h. All fluorescent images are 2D maximum intensity projections of 3D z-stacks generated using NIS-Elements (Nikon) software. For clear presentation of the hindbrain vasculature, the lateral dorsal aorta (located beneath the hindbrain vasculature in a dorsal view) was cropped out of the z-stack before 2D compression as previously described.^66^ Any image manipulation for brightness or contrast was equally applied to all images within an experiment and does not affect interpretation of data.

#### Survival analysis

To examine Il-1β-induced mortality, double transgenic *gfap:rtTA*, *TRE:Il1β^mat^* (i.e. CNS/Il-1β) embryos were induced with 0, 1, 2.5, 5, or 10 µg/mL Dox at approximately 6 hpf, and survival was monitored until 7 dpf. Survival assays were performed in 100×15 mm petri dishes (Falcon), and survival was tallied daily as dead embryos were identified and removed from the petri dishes. Dox was replaced daily with freshly prepared solution. Kaplan-Meier curves were made using Excel (Microsoft), and log rank tests were used to evaluate statistical significance.

#### Genome editing of zebrafish *il1r1* using CRISPR/Cas9

To rescue the Il-1β induced phenotypes in the CNS/Il-1β model, we performed genome editing of *il1r1* using a CRISPR/Cas9 strategy that utilized the crRNA:tracrRNA duplex format with recombinant *S. Pyogenes* Cas9 nuclease (Cas9) from Integrated DNA Technologies (IDT). Two CRISPR guide RNAs (crRNAs; cr1 and cr2) targeting the *il1r1* gene were previously designed as described by Sebo et al.^58^ The crRNA sequences are cr1, 5’-/AltR1/ucgacugcuggacaccagacguuuuagagcuaugcu/AltR2/-3’and cr2, 5’- /AltR1/uuaagguggagcuggucuuaguuuuagagcuaugcu/AltR2/-3’ against exon 8 and exon 9, respectively. CRISPR/Cas9 ribonucleoprotein (RNP) were prepared and microinjected into single-cell embryos according to the IDT demonstrated protocol “Zebrafish embryo microinjection” modified from Dr. Jeffrey Essner (Iowa State University) and as previously described. The embryos were treated with Dox and selected for no inflammation-related phenotypes. Those appearing healthy were raised to adulthood. To confirm the CRISPR-mediated deletion in both copies of *il1r1*, offspring were genotyped by extracting genomic DNA from individual embryos and performing PCR with *il1r1*-specific primers: forward primer 5’- tatgtgttcctcttgcagCG-3’ and reverse primer 5’-tgtttatacgagcacCTGTGG-3’ located at the intron 7/exon8 splice-acceptor site and the intron 9/exon 9 splice acceptor site, respectively (lower case denotes intron sequence and upper case denotes exon sequence).

The same cr1/cr2 RNP complexes were used to generate *il1r1-/-* crispants. Combined cr1/cr2 (1:1) RNP complexes were microinjected into CNS/Il-1β single-cell embryos and imaged by confocal microscopy at about 52-54 hpf to examine CtA formation using *kdrl:EGFP* and transcriptional activation of the *glut1b* using the transgenic line *glut1b:mCherry*. Images were quantified as described below.

#### Microangiography with fluorescent tracers

To examine permeability of the newly formed vessels, we microinjected fluorescent tracer molecules into circulation as previously described.^36^ CNS/Il-1β embryos with *kdrl:EGFP* were either untreated or treated with 1.0 μg/mL Dox at approximately 6 hpf. Embryos at 48-52 hpf were anesthetized in 0.02% tricaine (Western Chemical Inc.) and co-injected with approximately 4 nL of Texas Red dextran, 10kDa (Life Technologies) at 2 mg/mL and DAPI (Roche) at 4 mg/mL into the pericardial sac. At 25-35 minutes post-injection, embryos were embedded in 1.0% low-melting point agarose and imaged via confocal microscopy. For each embryo, the hindbrain was imaged followed immediately by imaging the central region of the trunk of the same embryo. Fluorescent signal was quantified as described below.

### QUANTIFICATION AND STATISTICAL ANALYSIS

#### Quantification of neutrophils and microglia/macrophages

The number of neutrophils (*mpx:EGFP*-positive cells) and microglia/macrophages (*mpeg1:mCherry*-positive cells) in the head at 4 dpf was quantified using FIJI (ImageJ) software. First, 2D maximum intensity projections (MIPs) were generated using NIS-Elements (Nikon) software. Using FIJI, these MIPs were converted to binary images using a minimum threshold of 15 for neutrophils and 20 for microglia/macrophages. The FIJI command “Watershed” was applied to the microglia/macrophages images to separate groups of cells; however, this was not required for the neutrophils. Finally, the number of cells for both types was counted using the FIJI command “Analyze Particles”. The minimum countable particle size was set to 50 pixels for neutrophils and 20 pixels for microglia/macrophages (pixel sizes were different). Statistics (p-values) for both groups were determined by one-tailed student t-test (*p < 0.05; **p < 0.01; ***p < 0.001; ns = not significant).

At earlier timepoints, both neutrophils (*mpx:EGFP*-positive cells) and microglia/macrophages (*mpeg1:mCherry*-positive cells) were counted manually (using NIS-Elements 3D rendering). For both cell types, the region of interest was restricted to inside the brain (not on the surface of the head) and the depth of the imaging covered half of the brain. Statistics (p-values) were determined by ANOVA with Tukey HSD post hoc test (*p < 0.05; ns = not significant).

#### Quantification of vasculature phenotypes

Zebrafish vasculature phenotypes were quantified using NIS-Elements (Nikon) and FIJI (ImageJ) software. To quantify CNS angiogenesis, the number of central artery loops (CtAs) were counted in the hindbrain at 2 dpf using the 3D rendering in NIS-Elements. Vessels were counted as CtAs if they branched upward from the primordial hindbrain channels and connected (directly or indirectly) to the basilar artery. To quantify the intersegmental vessels (ISVs) in the trunk of embryos, the number of ISVs at 2 dpf was counted using the 3D rendering in NIS-Elements. Only full, normally formed ISVs were counted.

Expression of *glut1b:mCherry* in hindbrain vasculature was quantified as the average mCherry fluorescence intensity within blood vessels. Confocal z-stacks showing *glut1b:mCherry* and *kdrl:EGFP* in separate channels were first cropped to include just the hindbrain using NIS-Elements. Next, a binary mask of the vasculature was created in FIJI using the zebrafish vasculature quantification (ZVQ) program and workflow developed by Kugler et al.^95^ Specifically, z-stacks showing the *kdrl:EGFP* signal were converted to tiffs and run through the vessel enhancement and vessel segmentation portions of the ZVQ workflow. The parameter σ was set to 3.5 and the threshold for segmentation was set to 15-255. The resulting binary mask z-stacks were then used along with 3D ROI Manager, a FIJI plug-in from the 3D ImageJ Suite developed by Ollion et al., to create 3D ROIs (regions of interest) specifically encapsulating the vasculature in each frame of the z-stack.^96^ Finally, using the 3D ROI Manager, these 3D ROIs were applied to the corresponding z-stacks showing the *glut1b:mCherry* signal in the same fish, and the mean signal intensity within the ROI was quantified for each stack using the “Quantif 3D” command in the 3D ROI Manager.

Expression of *glut1b:mCherry* in hindbrain vasculature was also quantified as the fraction of hindbrain vasculature labeled by *glut1b:mCherry* using the ratio of the length of *glut1b:mCherry*-labeled vasculature to the length of *kdrl:EGFP*-labeled vasculature. The length of hindbrain vasculature for each transgene was calculated in FIJI using the ZVQ program and workflow.^95^ Specifically, z-stacks of either the *kdrl:EGFP* signal or the *glut1b:mCherry* signal were run through the vessel enhancement and vessel segmentation portions of the ZVQ workflow as described above. Then, the resulting binary mask z-stacks were skeletonized and measured using the vascular quantification portion of the ZVQ workflow.

To measure transcriptional activation of the Wnt/β-catenin signaling pathway in hindbrain vasculature, we used a transgenic Wnt/β-catenin transcriptional reporter line with a nuclear localization signal, *TCF:mCherry*. To quantify expression of the transgene, we counted the number of *TCF:mCherry*-positive nuclei within the hindbrain vasculature (using NIS-Elements 3D rendering). This number was then divided by the length of the hindbrain vasculature in that embryo to account for changes in the total number of endothelial cells. The length of hindbrain vasculature was calculated from z-stacks of the *kdrl:EGFP* signal in FIJI using the ZVQ program and workflow as described above for the fraction of *glut1b:mCherry*.^95^

Statistics (p-values) were determined by ANOVA with Tukey HSD post hoc test for most experiments and by two-tailed student t-test for ISVs and controls (*p < 0.05; **p < 0.01; ***p < 0.001; ns = not significant).

#### Quantification of fluorescent tracers

The leakage of the Texas Red dextran (10 kDa) tracer from the hindbrain vasculature was quantified using relative fluorescence intensity. First, a binary mask of the vasculature (as labeled by *kdrl:EGFP*) was created in FIJI using the vessel segmentation portions of the ZVQ workflow and the masks were used to create 3D ROIs as described for the *glut1b:mCherry* quantification above.^95, 96^ These 3D ROIs were applied to z-stacks showing the Texas Red signal and all signal was deleted from inside the ROI (inside the vasculature) using the “Erase” command in the 3D ROI Manager. Next, 3D ROIs were created to limit the region of interest to only the area around the CtAs in the center of the hindbrain channel (see dashed boxes in Supplemental Figure 4A). The mean signal intensity of the Texas Red within these ROIs was then quantified using the “Quantif 3D” command in the 3D ROI Manager. Finally, these intensity values were normalized by the mean fluorescence intensity of Texas Red within the dorsal aorta. This was quantified for each fish from the corresponding tail image in FIJI. First, the *kdrl:EGFP* signal was used to outline an ROI within the dorsal aorta and spanning two segments of the trunk. Then the mean signal intensity of Texas Red in the region was measured. Statistics (p-values) were determined by one-tailed student t-test (ns = not significant).

### KEY RESOURCES TABLE

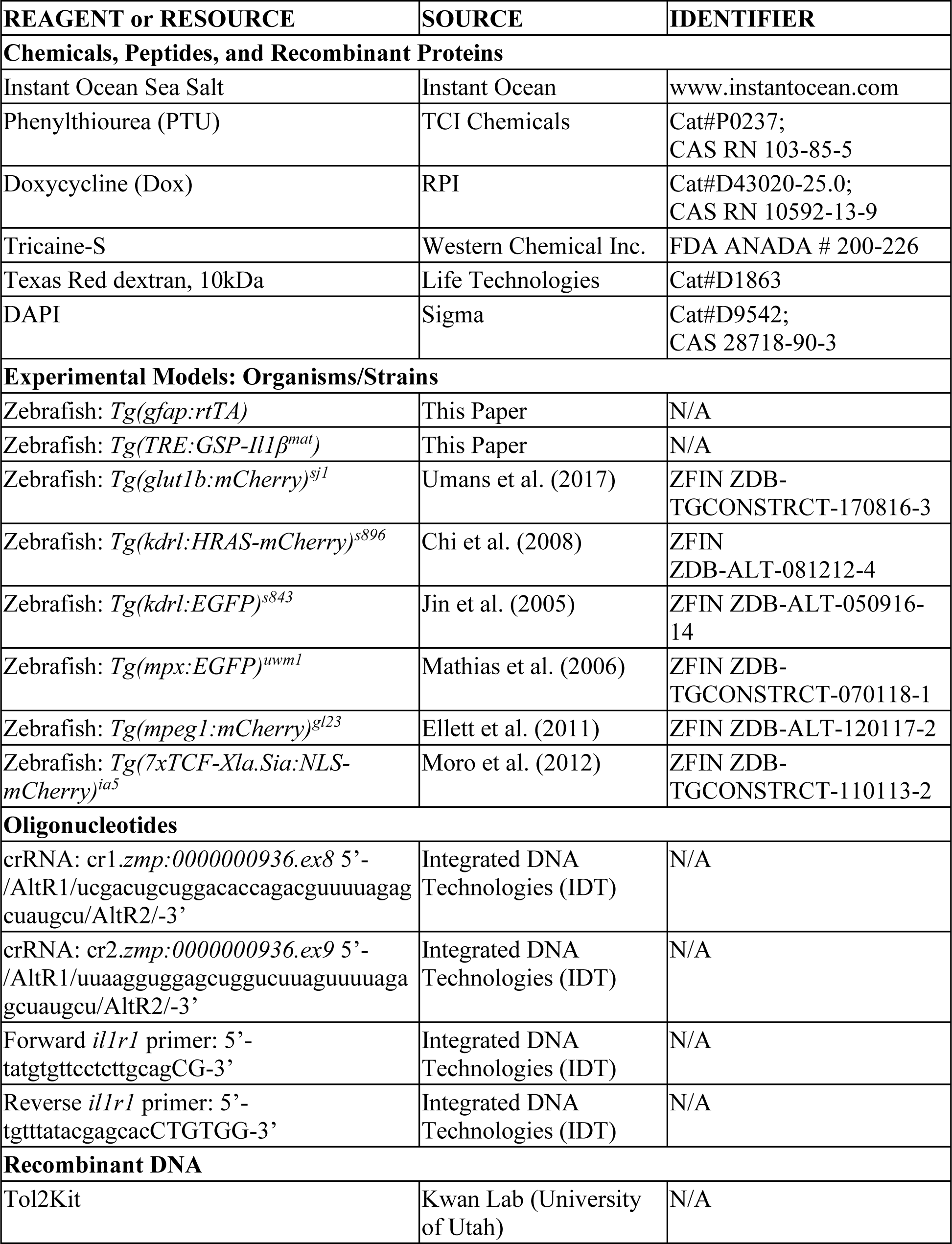
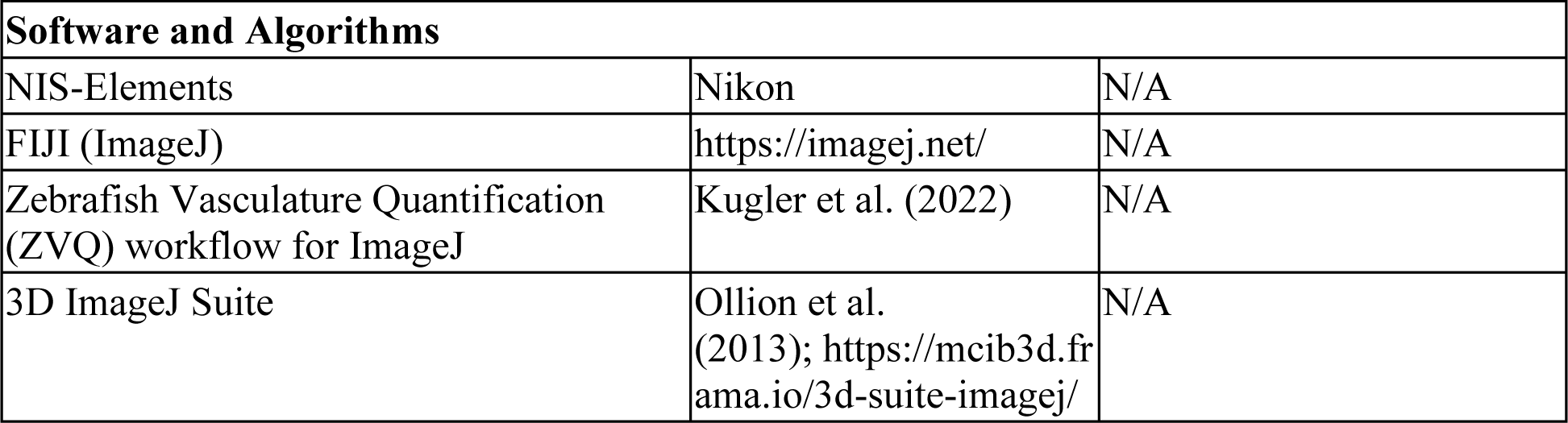

## Supplemental Figures and Legends

**Figure S1.**
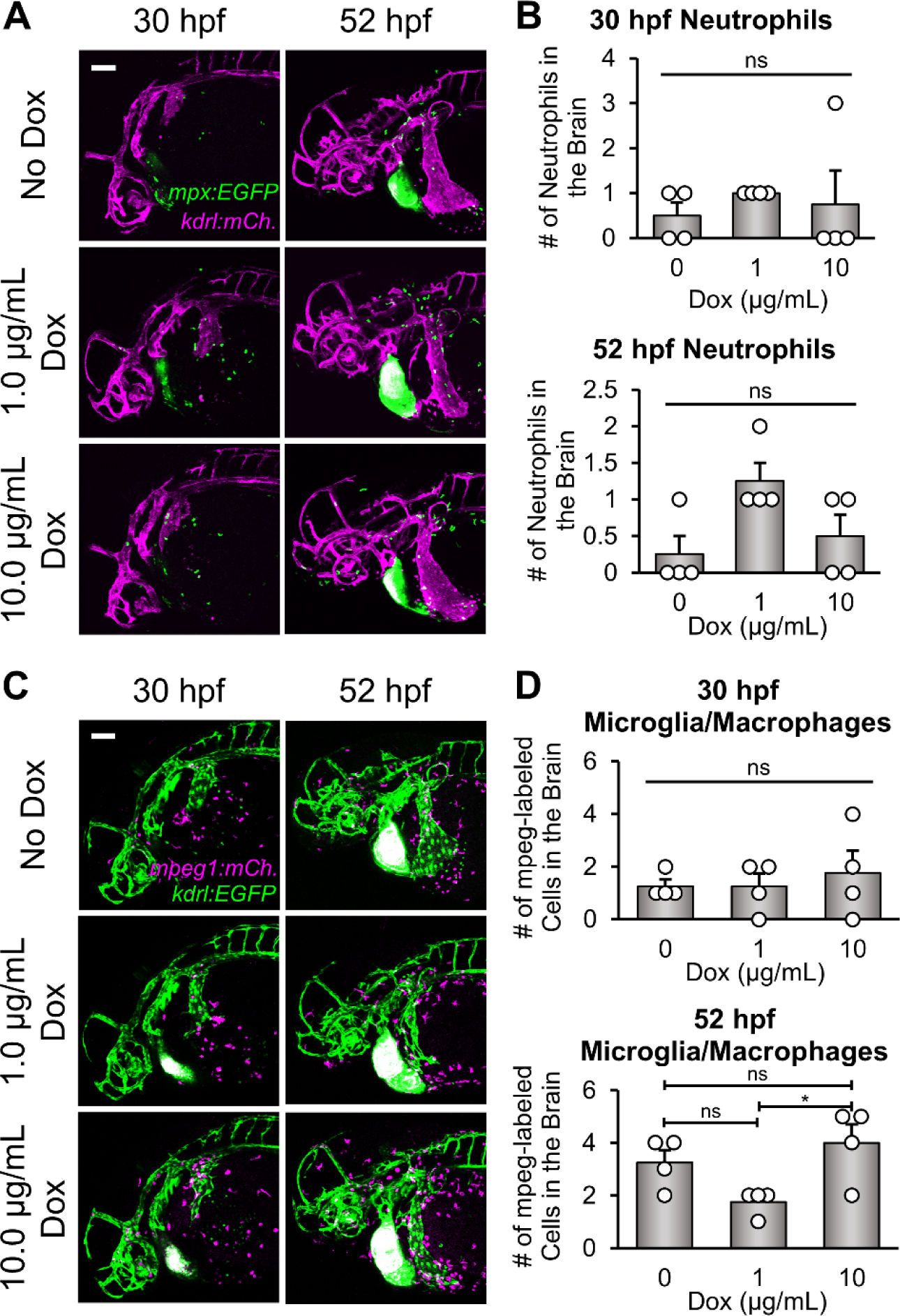
Embryos at 30 and 52 hpf show low numbers of neutrophils and microglia/macrophages in the brain regardless of Il-1β expression. (**A**) Representative confocal microscopy images showing neutrophils (*mpx:EGFP*; green) and blood vessels (*kdrl:mCherry*; magenta). CNS/Il-1β embryos were untreated (No Dox) or treated with 1.0 or 10.0 μg/mL of Dox at 6 hpf and then imaged at 30 or 52 hpf (lateral view; dorsal top; anterior left). Scale bar is 100 μm. (**B**) Quantification of neutrophils (*mpx:EGFP*) in the brains of untreated (No Dox) or treated (1.0 or 10.0 μg/mL Dox) CNS/Il-1β embryos at 30 or 52 hpf (*n=*4). (**C**) Representative confocal microscopy images showing microglia/macrophages (*mpeg1:mCherry*; magenta) and blood vessels (*kdrl:EGFP*; green). CNS/Il-1β embryos were untreated (No Dox) or treated (1.0 or 10.0 μg/mL Dox) at 6 hpf and then imaged at 30 or 52 hpf (lateral view; dorsal top; anterior left). Scale bar is 100 μm. (**D**) Quantification of microglia/macrophages (*mpeg1:mCherry*) in the brains of untreated (No Dox) or treated (1.0 or 10.0 μg/mL Dox) CNS/Il-1β embryos at 30 or 52 hpf (*n=*4). Error bars in B and D represent means ± SEM (*p < 0.05; ns = not significant).

**Figure S2.**
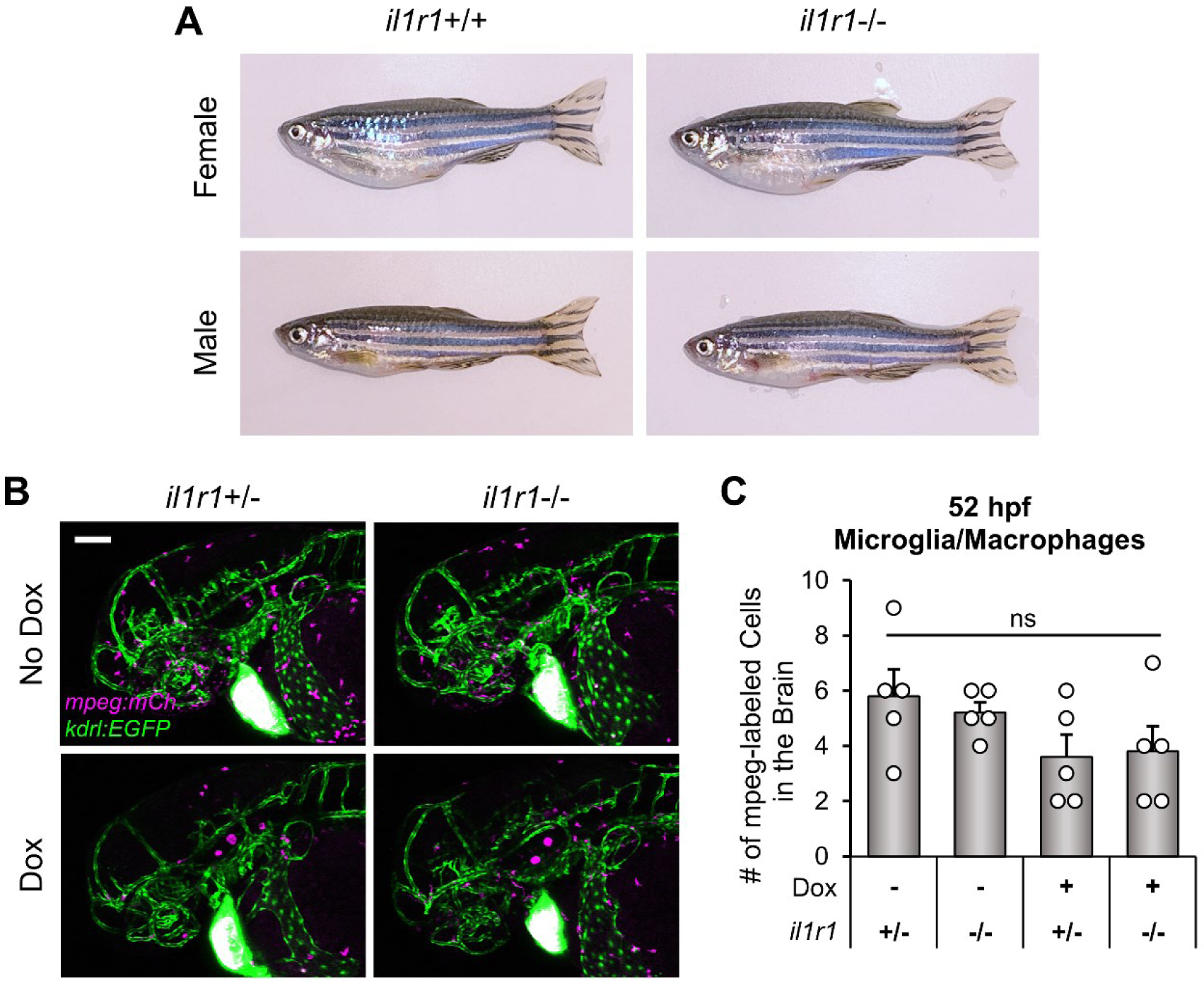
*il1r1-/-* zebrafish show no differences in adult gross morphology or embryonic microglia/macrophage cell populations. (**A**) Representative digital images showing adult CNS/Il-1β, *kdrl:EGFP* zebrafish that were either *il1r1+/+* or *il1r1-/-* (lateral view; dorsal top; anterior left). (**B**) Representative confocal microscopy images showing microglia/macrophages (*mpeg1:mCherry*; magenta) and blood vessels (*kdrl:EGFP*; green) in the head region. CNS/Il-1β *il1r1+/-* or *il1r1-/-* embryos were either untreated (No Dox) or treated (10.0 μg/mL Dox) at 6 hpf and then imaged at 52 hpf (lateral view; dorsal top; anterior left). Scale bar is 100 μm. (**C**) Quantification of microglia/macrophages (*mpeg1:mCherry*) in the brains of untreated (No Dox) or treated (10.0 μg/mL Dox) CNS/Il-1β *il1r1+/-* or *il1r1-/-* embryos at 52 hpf (*n=*5). Error bars represent means ± SEM (ns = not significant).

**Figure S3.**
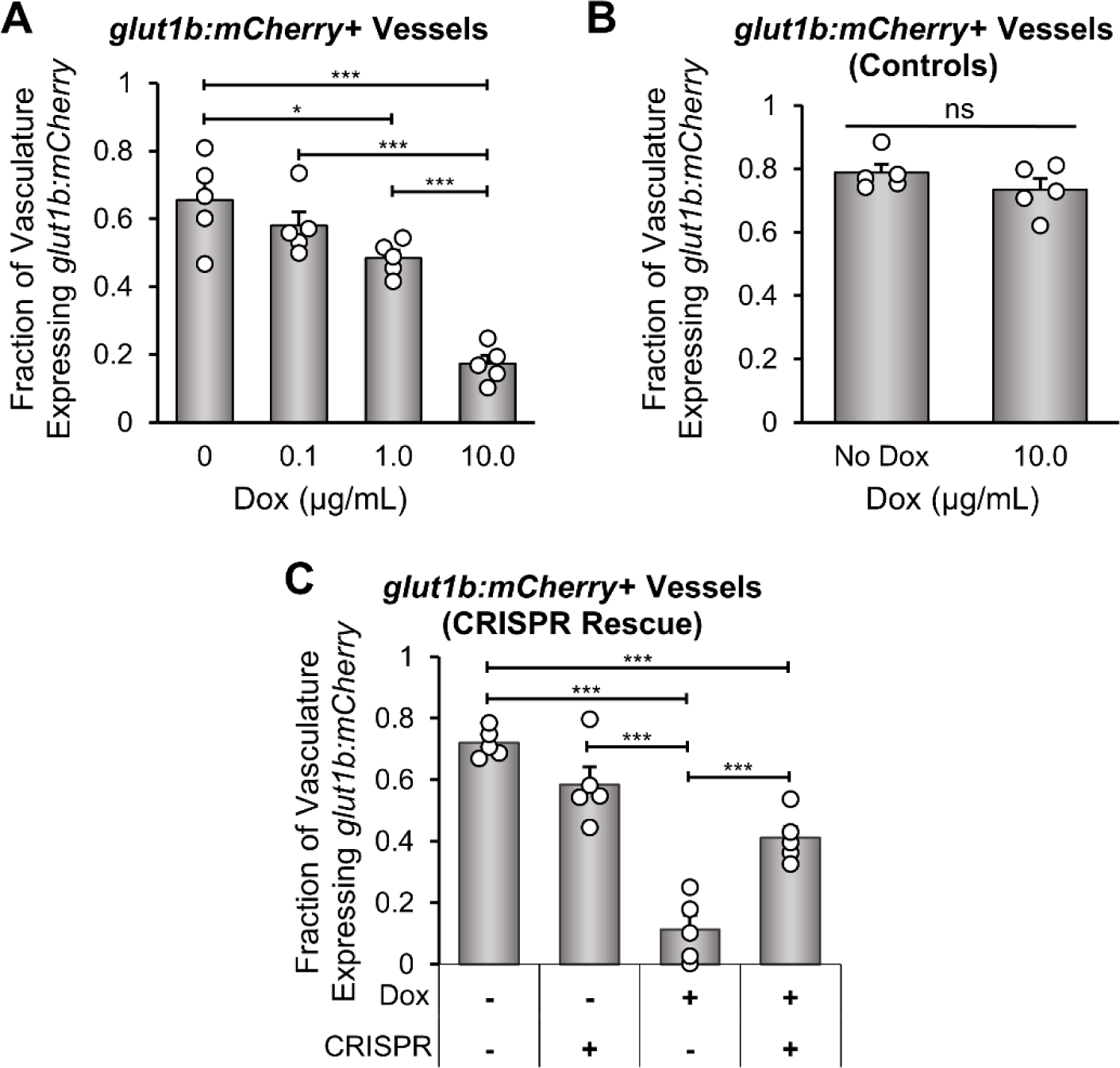
Fraction of *kdrl:EGFP*-labeled hindbrain vasculature expressing *glut1b:mCherry*. (**A**) Quantification of the fraction of hindbrain vasculature expressing *glut1b:mCherry* (length of *glut1b:mCherry* vasculature/length of *kdrl:EGFP* vasculature) at 52 hpf in untreated (No Dox) or treated (0.1, 1.0, or 10.0 μg/mL Dox) CNS/Il-1β embryos (*n=*5). (**B**) Quantification of the fraction of hindbrain vasculature expressing *glut1b:mCherry* (length of *glut1b:mCherry* vasculature/length of *kdrl:EGFP* vasculature) at 52 hpf in control *kdrl:EGFP*, *glut1b:mCherry* embryos (no CNS/Il-1β) without (No Dox) or with (10.0 μg/mL Dox) treatment (*n=*5). (**C**) Quantification of the fraction of hindbrain vasculature expressing *glut1b:mCherry* (length of *glut1b:mCherry* vasculature/length of *kdrl:EGFP* vasculature) at 52 hpf in untreated (No Dox) or treated (10.0 μg/mL Dox) CNS/Il-1β embryos with or without *il1r1*-targeted CRISPR injection (*n=*5). Note that these data follow the same trend as the average *glut1b:mCherry* fluorescence intensity in hindbrain vasculature. Error bars in all panels represent means ± SEM (*p < 0.05; ***p < 0.001; ns or no label = not significant).

**Figure S4.**
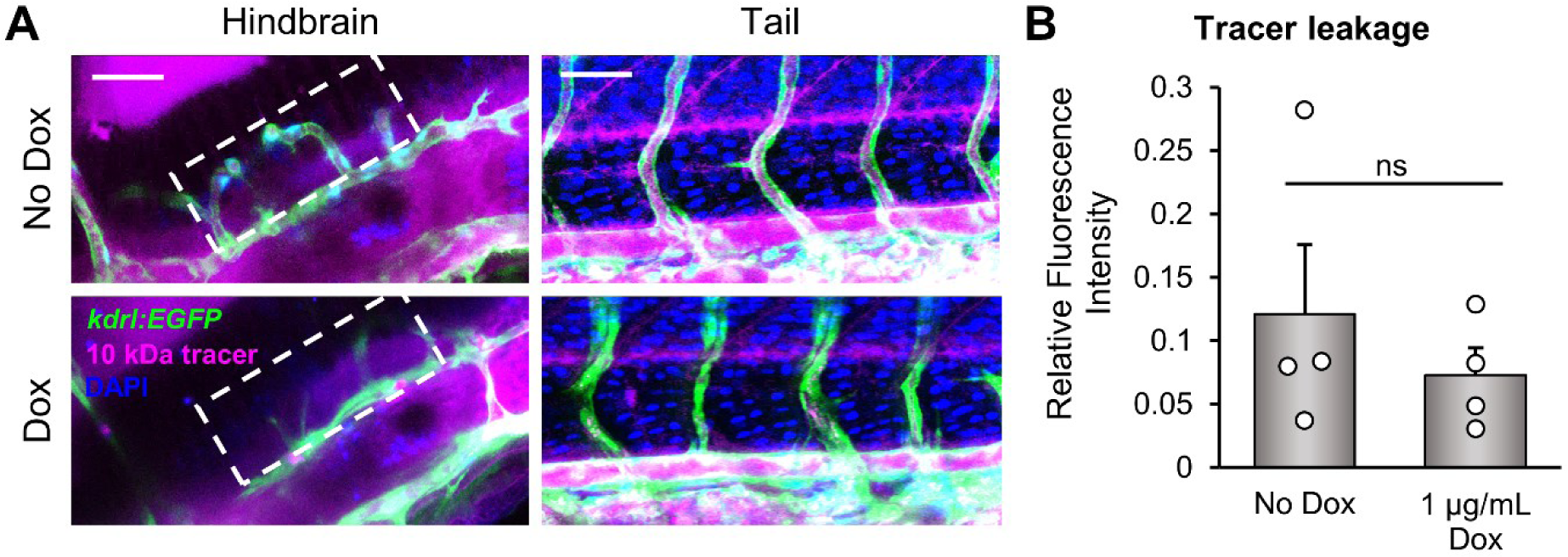
Dox-induced Il-1β expression does not affect leakage of injected tracers at 2 dpf. (**A**) Representative confocal microscopy images showing *kdrl:EGFP* (green) CNS/Il-1β embryos that were either untreated (No Dox) or treated (1.0 μg/mL Dox) at 6 hpf and then injected in the heart with DAPI (blue) and Texas Red dextran, 10 kDa (magenta) at 48 hpf. Images (lateral view; dorsal top; anterior left) show the hindbrain (left) and the center of the tail (right) 30 minutes after injection. Scale bars are 50 μm. (**B**) Quantification of leakage of Texas Red dextran, 10 kDa (magenta) from the hindbrain vasculature (region shown by dashed boxes in panel A) of untreated (No Dox) or treated (1.0 μg/mL Dox) CNS/Il-1β embryos 30 minutes after injection normalized by the fluorescent intensity within the dorsal aorta (*n=*4). Error bars represent means ± SEM (ns = not significant).

